# High-dose rifamycins enable shorter oral treatment in a murine model of *Mycobacterium ulcerans* disease

**DOI:** 10.1101/395095

**Authors:** Till F. Omansen, Deepak Almeida, Paul J. Converse, Si-Yang Li, Jin Lee, Ymkje Stienstra, Tjip van der Werf, Jacques H. Grosset, Eric L. Nuermberger

**Affiliations:** Center for Tuberculosis Research, Department of Medicine, Johns Hopkins University, Baltimore, Maryland, USA; Infectious Diseases Unit, Department of Internal Medicine, University of Groningen, Groningen, The Netherlands; Department of Pulmonary Diseases and Tuberculosis, University of Groningen, Groningen, The Netherlands

**Author notes:** Address correspondence to: Dr. Eric Nuermberger.

## Abstract

Buruli ulcer (BU), caused by *Mycobacterium ulcerans,* is a neglected tropical skin and soft tissue infection that is associated with disability and social stigma. The mainstay of BU treatment is an eight-week course of 10 mg/kg rifampin (RIF) and 150 mg/kg streptomycin (STR). Recently, the injectable STR has been shown to be replaceable with oral clarithromycin (CLR) for smaller lesions for the last four weeks of treatment. A shorter, all-oral, highly efficient regimen for BU is needed, as the long treatment duration and indirect costs currently burden patients and health systems. Increasing the dose of RIF or replacing it with the more potent rifamycin drug rifapentine (RPT) could provide such a regimen. Here, we performed a dose-ranging experiment of RIF and RPT in combination with CLR over four weeks of treatment in a mouse model of *M. ulcerans* disease. A clear dose-dependent effect of RIF on both clinical and microbiological outcomes was found, with no ceiling effect observed with tested doses up to 40 mg/kg. RPT-containing regimens were more effective on *M. ulcerans*. All RPT-containing regimens achieved culture negativity after only four weeks while only the regimen with the highest RIF dose (40 mg/kg) did so. We conclude that there is dose-dependent efficacy of both RIF and RPT and that a ceiling effect is not reached with the current standard regimen used in the clinic. A regimen based on higher rifamycin doses that are currently being evaluated against tuberculosis in clinical trials could shorten and improve therapy of Buruli ulcer.

## Introduction

Buruli ulcer (BU) is a neglected tropical disease caused by *Mycobacterium ulcerans*. This pathogen is phylogenetically related to *M. tuberculosis* and *M. leprae* and is believed to have evolved from a common ancestor, *M. marinum* (1). BU presents as skin and soft tissue lesions, potentially leading to scarring, contracture, physical impairment and psychosocial exclusion of patients due to disfiguring wounds (2). In 2015, BU was reported from 33 countries mainly in sub-Saharan West Africa, where it predominantly occurs in rural communities (3). BU treatment formerly consisted of wide surgical excision and skin grafting (4). Following a phase of antibiotic regimen discovery and development in a murine model of BU and a subsequent clinical trial, the World Health Organization (WHO) recommended treatment with rifampin (RIF) 10 mg/kg body weight orally and streptomycin (STR) 15 mg/kg by intramuscular injection, both administered daily for 8 weeks. Despite good clinical outcomes with this regimen (5, 6), injectable streptomycin causes ototoxicity in 25-30% of patients, nephrotoxicity and risk of infection by contaminated needles (7). Streptomycin can be replaced by clarithromycin (CLR), which has a largely bacteriostatic effect on *M. ulcerans*, for the last four weeks of the eight-week regimen for early, limited lesions (8). An ongoing trial (ClinicalTrials.gov Identifier: NCT01659437) is investigating an all-oral eight-week RIF+CLR regimen with the standard dose of RIF 10 mg/kg. Still, despite free treatment in most countries, the indirect costs of hospitalization and loss of income burden patients and families (9, 10). A shortened, highly effective, all-oral regimen, is urgently needed to improve care for this neglected tropical disease.

Rifampin is a rifamycin antibiotic and the mainstay oral drug in treatment of *M. ulcerans*. The currently recommended dose of 10 mg/kg has been adopted from regimens to treat tuberculosis (TB), where this dose was chosen mainly for economic reasons and in fear of toxicity associated with high-dose intermittent regimens when the drug was first introduced in the 1970’s (11). Recently, interest in using higher doses of RIF as a means to shorten the duration of treatment for TB has intensified. Trials have demonstrated that treatment with up to 35 mg/kg RIF appears to be safe and offers a nonlinear increase in exposure to the drug that is associated with more rapid sputum culture conversion (12, 13). Further studies showed that high-dose RIF-containing regimens improve and shorten the duration of TB therapy (14-16). Rifapentine (RPT) is another rifamycin antibiotic that is active against *M. tuberculosis* and *M. ulcerans*. It is slightly more active *in vitro* against *M. ulcerans*, with an MIC of 0.125 µg/ml compared to RIF (0.25 µg/ml) and is effective *in vivo* in mice at a daily dose of 10 mg/kg (17). Its longer half-life also makes it an attractive alternative to RIF. Higher daily doses of RPT are more active in murine models of TB (18). Furthermore, RPT doses up to 20 mg/kg were safe in humans (19) and shown to increase the rate of sputum culture conversion in TB patients in an exposure-dependent manner that may lead to shorter treatment durations for TB (20, 21). In Buruli ulcer, a regimen of RPT at 10 mg/kg/day plus CLR was shown to be more effective than RIF+STR in a mice (22). Also, combining RIF at 10 to 40 mg/kg/day with clofazimine were shown to reduce treatment duration in the Buruli ulcer footpad model, recently (23). We hypothesize that regimens containing high-dose RIF or RPT will be a next pivotal step in improving Buruli ulcer chemotherapy by increasing efficacy and reducing duration.

To evaluate an all-oral regimen with high-dose RIF or RPT in combination with the standard macrolide used today, CLR, we performed dose-ranging experiments with escalating doses of both drugs in combination with CLR in the mouse footpad model of BU. We identified high-dose rifamycin-containing regimens that are more bactericidal than the same all-oral regimen with RIF at the standard dose. These regimens can be evaluated in clinical trials in hopes of unburdening patients from long-term treatment, reducing indirect costs and improving adherence.

## Materials and methods

### Ethical clearance

Animal experiments described in this study were conducted in strict adherence with the Animal Welfare Act and Public Health Service Policy. Experiments were performed at the Johns Hopkins University which is accredited by the private Association for the Assessment and Accreditation of Laboratory Animal Care International. All procedures involving mice were approved by the Johns Hopkins University Animal Care and Use Committee.

### Bacteria

*M. ulcerans* 1059 is an isolate originating from a clinical specimen from a patient in Ghana. An autoluminescent version of this isolate, Mu1059AL, was generated in our laboratory, as previously described (24, 25). Bacteria were passaged in mouse footpads and frozen footpad suspensions were used for infection of mice in this study. To quantify colony forming units (CFU), samples of bacterial cultures and footpad homogenates were cultured in serial 10-fold dilutions on Middlebrook 7H11 selective agar (Becton-Dickinson, Sparks, MD) at 32°C.

### Antibiotics

RIF and STR were purchased from Sigma (St. Louis, MO, USA). RPT and CLR were kindly provided by Sanofi (Bridgewater, NJ, USA) and Abbott (Abbott Park, IL, USA), respectively. RIF, RPT and STR were dissolved in distilled water, while CLR was suspended in distilled water with 0.05% agarose. The doses for RIF were 5, 10, 20, and 40 mg/kg; the doses for RPT were 5, 10, and 20 mg/kg; the dose for STR was 150 mg/kg, and the dose for CLR was 100 mg/kg. The doses for the rifamycins produce exposures similar to the average plasma AUC observed in humans at doses up to 35 mg/kg for RIF and 20 mg/kg for RPT (16, 26). The doses for STR and CLR produce plasma AUC values similar to human doses of 15 mg/kg and 7.5 mg/kg, respectively (27-30).

### Infection and treatment

Female BALB/c mice (n=110), aged 4-6 weeks were purchased from Charles River (Wilmington, MA, USA) and allowed to acclimatize for 5 days upon arrival at the facility. Food and water were provided *ad libitum*. Mice were infected with approximately 4.56 log_10_ CFU of Mu1059AL in 0.03 ml PBS into both hind footpads via subcutaneous injection. Five untreated mice were sacrificed the following day to confirm the number of implanted CFU. Animals were regularly checked for signs of general illness and progression of footpad infection. The swelling grade was used to evaluate *M. ulcerans*-induced footpad pathology, as previously described (29). A swelling grade of 0 corresponds to a normal footpad, grade 1 to a non-inflammatory but swollen footpad, grade 2 to inflammatory swelling with edema and redness and 3 swelling of the entire hind-foot. At week 6 post-infection (day 0), animals were randomized into treatment groups and treatment was initiated. Drugs were administered once daily, 5 days per week. STR was administered by subcutaneous injection. All other drugs were administered in 0.2 ml via esophageal gavage. Sacrifices for CFU counts were always performed three days after the last dose of antibiotic was administered to minimize drug carryover effect.

### Treatment outcome monitoring

After treatment commenced, the swelling grade and relative light units (RLU) emitted from autoluminescent bacteria in the footpads were measured weekly, as previously described (25), while mice were anesthetized via intraperitoneal injection of ketamine/xylazine 87.5/12.5 mg/kg (Zoetis, Kalamazoo, MI). At day 0, week 2 and week 4, 5 mice per group were sacrificed to assess CFU counts. Mouse footpad tissue was carefully harvested from both hind footpads with the exception of the well-defined control groups R10, R10C100 and R10C150 where only one footpad was analyzed per mouse. Tissue was finely minced with scissors before being suspended in 2.0 ml PBS. The suspensions were vortexed for homogenization and quantitative bacterial cultures were performed as described above. Half of the footpad homogenate was plated and the lower limit of detection was 2 CFU/footpad.

### Data analysis

CFU counts were log-transformed. To compare mean CFU between treatment groups, student’s t-test and analysis of variance (ANOVA) were used. The value of α was set at 0.05. Data were analyzed and graphs were computed using GraphPad Prism version 7.0a (GraphPad Software, Inc., San Diego, CA).

## Results

### Infection

Successful implantation of viable bacteria was confirmed by a mean (±SD) footpad CFU count of 4.09 log_10_ (± 0.23) CFU on the day after infection. In untreated animals, clinical pathology, as assessed by swelling grade, stayed at a level of 2 during the first two weeks of the treatment phase and improved to 1.25 in week 4 (Fig 1). RLU also declined over time in untreated mice, however less so than in treated animals (Fig 2).

**Figure 1:**
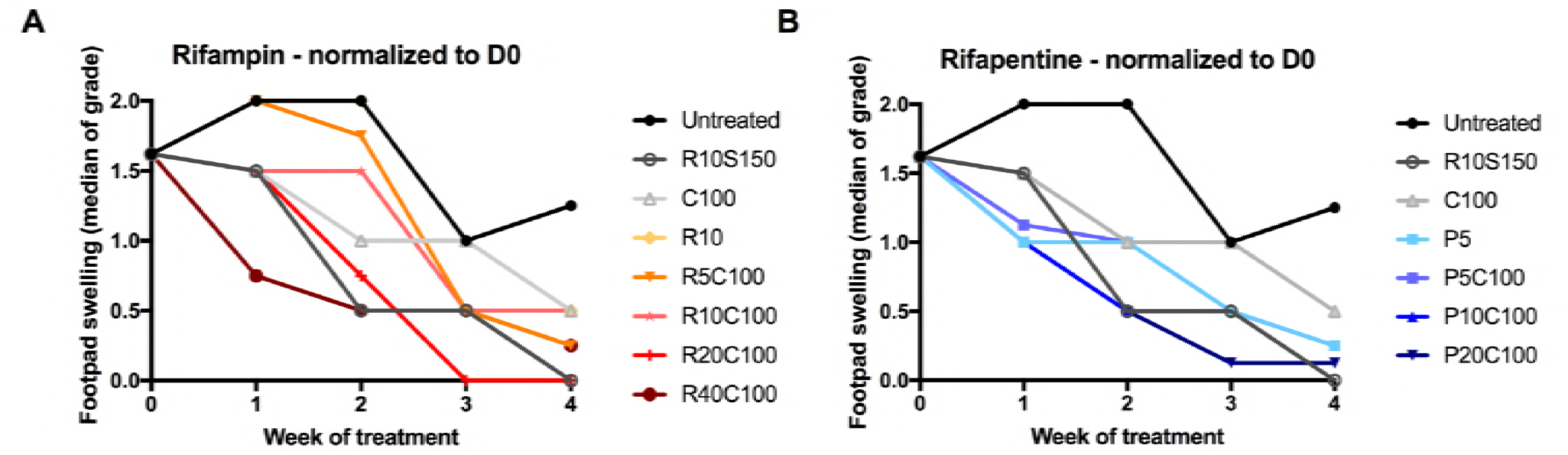
Median of footpad swelling grade of infected mouse footpads in response to treatment with high-dose rifamycins and clarithromycin. Treatment was initiated 6 weeks after infection, when swelling approached swelling grade 2. Swelling grade = 0 corresponds to no clinically visible pathology. Swelling grade = 1 infers redness of the footpad, grade = 2 edematous swelling of the footpad and grade = 3 ascending swelling of the leg and impeding necrosis. Data points represent medians per treatment group. Data were normalized to day 0 (beginning of treatment) by subtracting from the median swelling grade of all mice at D0 and assuming the total median as group mean for that time-point. All regimens reduced swelling grade compared to untreated mice. There was a visible dose-dependent effect; with escalating doses of rifampin (A) and rifapentine (B), swelling grade was reduced more drastically. Minimum swelling grade values of 0.23 and 0.15 were recorded at week 4 for the highest-dose regimens R40C100 and P20C100 respectively. D, day, R, Rifampin; S, Streptomycin; C, Clarithromycin; P, Rifapentine, numbers after drugs indicate doses in mg/kg.

**Figure 2:**
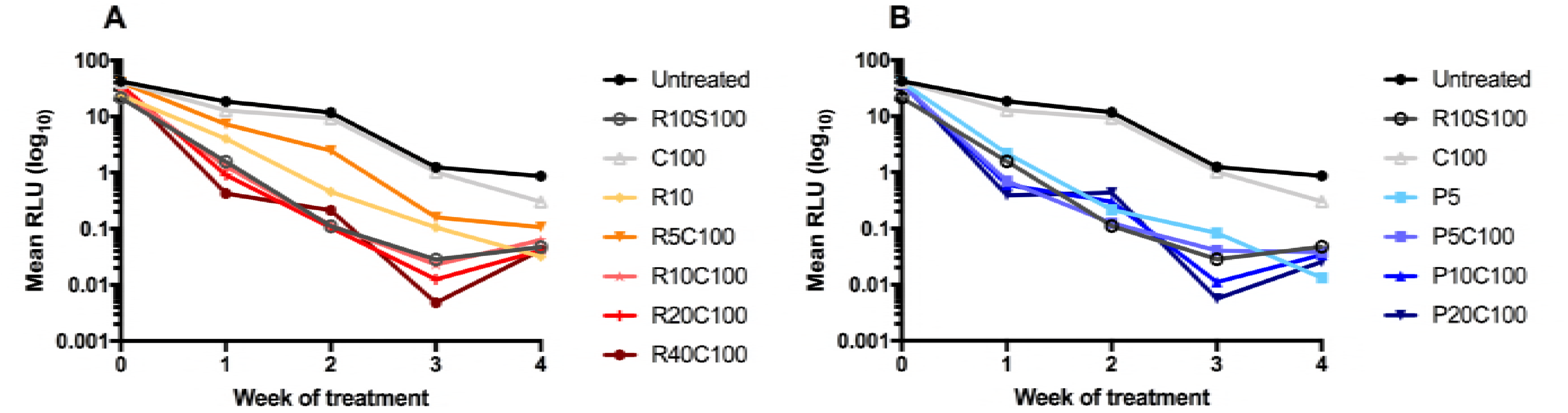
Mean relative light unit (RLU) counts from infected mouse footpads in response to treatment with escalating doses of rifamycins. Mice were infected with an autoluminescent strain of *M. ulcerans* that emits light when metabolically active. The mean per group RLU values are displayed, compared to Day 0, when the mean RLU was 35.09 (± 6.89). As observed in previous experiments, RLU counts stagnate at 7-9 weeks post infection (i.e. here 3 weeks of treatment), when a static bacterial growth phase is reached, as seen in untreated mice. Treatment with rifamycins radically reduced RLU counts compared to untreated and CLR-only treated mice. R, Rifampin; S, Streptomycin; C, Clarithromycin; P, Rifapentine.

### Response to treatment

Other than C100, all treatment regimens reduced the number of RLU measured from footpads. R5C100 and R10 were somewhat less effective than the other combination treatments which all had very similar effects on RLU counts. The swelling grade, however was affected in a more dose-dependent manner. At week 2, median swelling of untreated mice was 2 and CFU log_10_ 6.31 (± 0.31). The positive control regimen R10S150 led to rapid clinical improvement as shown by decreased swelling grade (median 0.5), which is in agreement with previous studies (31). Monotherapy with CLR100 performed worst in all parameters, as expected. All test regimens resulted in a statistically significant (p<0.0015 – 0.0001) reduction in CFU by week 2, compared to untreated mice, except RIF10 and R5C100 (Fig 3, Table S1). Animals treated with C100 and P5 were not sacrificed for CFU evaluation at week 2. RIF-containing combination regimens efficiently reduced footpad CFU counts by week 2 in a dose dependent fashion (Fig 3). RPT-containing regimens had a larger effect in terms of swelling grade and CFU reduction than RIF-containing regimens (Fig 3). All groups receiving rifapentine or rifampin at ≥ 20 mg/kg had mean CFU counts as low or lower than the rifampin-streptomycin control. Although no dose-response effect was observed for RPT, the mean CFU count in the P5C100-treated group includes one footpad with no detectable CFU which was likely due to a non-productive infection. Excluding this footpad, the mean CFU count is 2.98±0.96. After 4 weeks of treatment, swelling grade was reduced to a minimum median of 0.25 and 0.125 in the high-dose R40C100 and P20C100 groups, respectively. In terms of CFU reduction, all treatment groups, except C100, resulted in a statistically significant reduction of CFU, when compared to untreated mice (p=0.030 – 0.001). All high-dose RIF- and RPT-containing regimens performed better than standard R10C100 (p=0.001) and no worse than the RS control. Low-dose R5C100 performed less well, especially if 2 footpads with no detectable CFU were excluded as possibly never productively infected (in which case the mean CFU count was 2.64±1.09. Footpads from the R40C100 group and all RPT-containing regimens were culture-negative after 4 weeks of treatment (i.e. no colonies could be observed after plating the undiluted footpad homogenate). The CFU outcome of R10 and R10CLR100 after four weeks of treatment were both 1.77 (± 0.39 – 0.44). The addition of CLR to R10 thus seemed to have no effect on the outcome in this experiment.

**Figure 3:**
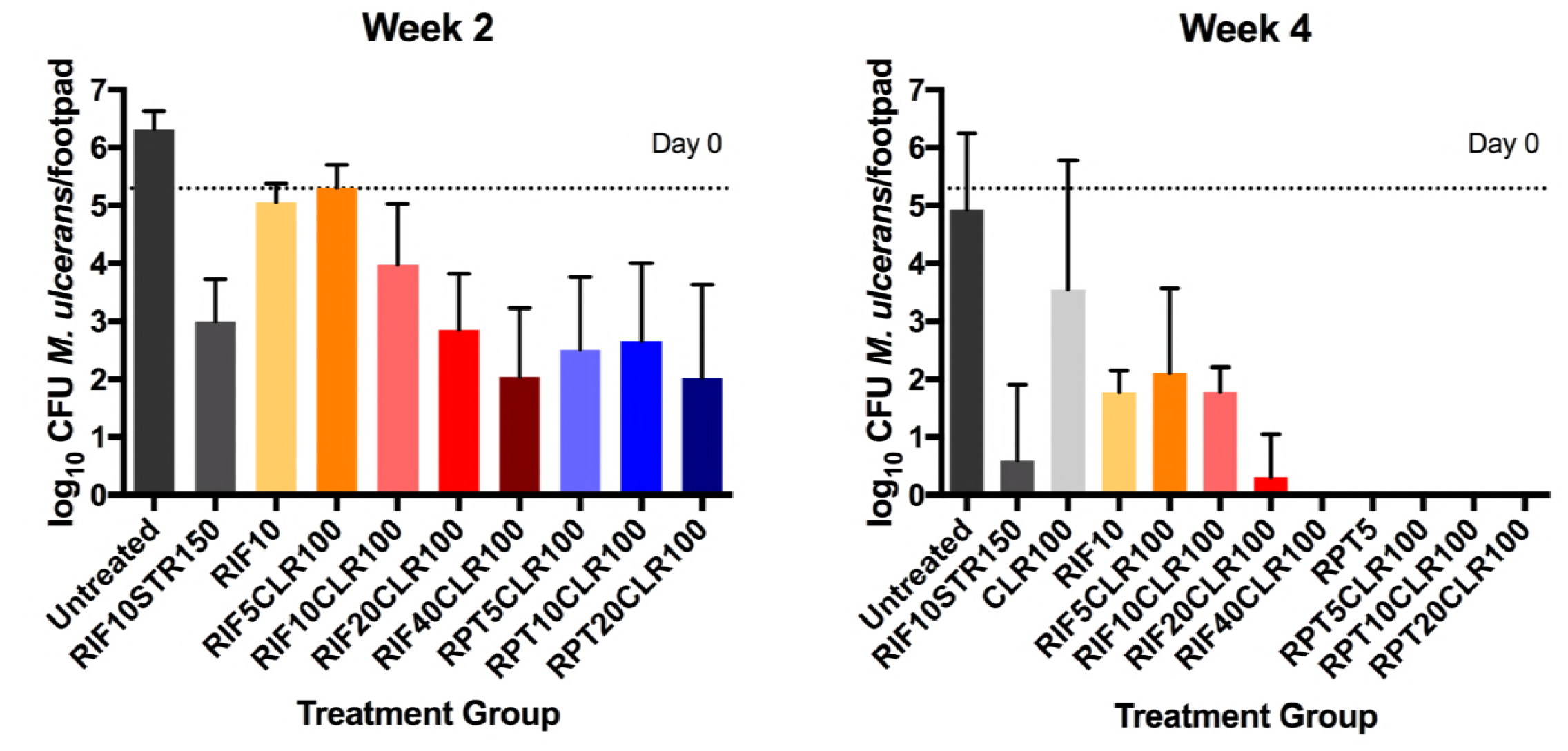
Microbiological outcome after two (A) and four (B) weeks of treatment with rifamycin-containing regimens. Mice were infected with 4.56 log10 colony forming units (CFU) of *M. ulcerans* into hind footpads. After 6 weeks of incubation, treatment was initiated (D0). At this time-point, CFU equaled 5.77 (± 0.60). Groups of mice (n=5) were sacrificed at week 2 and week 4, footpads dissected and minced and plated on 7H11 agar for colony counting and CFU analysis. There was a dose dependent reduction in CFU with elevating rifamycin doses. No ceiling effect was observed for R. At week 4, all rifapentine (P) containing regimens were culture negative, as was R40C100. CFU, colony forming unit; R, Rifampin; S, Streptomycin; C, Clarithromycin; P, Rifapentine, numbers after drugs indicate doses in mg/kg.

The previously WHO recommended treatment of R10S150 resulted in the highest healing rate, as evidenced by the finding that 80% of treated animals exhibited a swelling grade of zero upon completion of treatment. The swelling grade was reduced to zero in 60% and 50% of animals receiving R20C100 and P20C100, respectively.

## Discussion

Drug development for Buruli ulcer is greatly hampered by little economic interest for this neglected tropical disease. Repurposing and refining existing antibiotic regimens is a viable, rapid and relatively cost-effective measure to provide better care for BU patients. Here, we re-evaluated the dosage of RIF in the RC regimen for BU where the combination of RIF 40 mg/kg with CLR 100 mg/kg, as well as all RPT-containing regimens were most efficacious in reducing BU disease, as measured by swelling grade, and hastening the time to culture-negativity in the mouse footpads. These findings are timely and important because high-dose RIF- and RPT- containing regimens are currently being evaluated in phase 3 clinical trials for the ability to shorten the duration of TB treatment after promising results in phase 2 trials (16, 20). Our results indicate that similar dose optimization should be explored to enhance the efficiency of all-oral regimens for BU.

Shorter treatment would benefit patients by reducing indirect costs and barriers to treatment, and increasing adherence (9, 10, 32). With RIF, a clear dose-response relationship was demonstrated by linear regression with both clinical (swelling grade) and microbiological (CFU) outcome. There appeared to be no ceiling effect with the doses tested. In contrast, for RPT, no dose-response effect was evident at week 2 and the CFU counts were reduced to zero in all groups receiving RPT at week 4 of treatment. It is possible that the pharmacological ceiling was reached. However, this would be surprising given the increasing dose-response relationship up to RPT doses of 160 mg/kg that we previously observed in a mouse model of TB (33). Despite a three-day interval between the last dose of drug and sacrificing animals for microbiological analysis of footpads, we cannot exclude RPT carryover owing to its long half-life and possible elevated tissue concentration (17).

A dose-response relationship for RPT may have been evident if we had held mice beyond the end of treatment to determine if relapse occurred. While demonstrating cure without relapse is considered to be the gold standard outcome for pre-clinical TB efficacy models (34), culture negativity and relapse-free follow-up rates are debated as outcomes of choice in antimicrobial evaluation for *M. ulcerans* infection models (17). The hallmark of *M. ulcerans* infection is the presence of mycolactone, the analgesic, necrosis-causing and immune-suppressive toxin (35, 36). Reduction of the bacterial burden to a critical threshold at which the host immune response gains foothold enough to clear the remainder of bacteria may be sufficient to successfully treat the infection. If so, then comparing regimens on the basis of bactericidal activity alone, and not a relapse endpoint, would be reasonable. Relapse after successful antimicrobial therapy of *M. ulcerans* infection is exceedingly rare. For example, Klis et al. have shown that defaulters from the 8-week regimen often proceeded to heal their lesions despite the early discontinuation of antimicrobial therapy (37). We hypothesize that more mice from culture-negative as well as low-CFU treatment groups would have progressed to swelling grade 0 during follow-up.

Integrated approaches to control neglected tropical diseases are increasingly advocated. It is thus noteworthy that high-dose, short-course regimens with rifampicin are being investigated as potential therapeutic agents for lymphatic filariasis and onchocerciasis (38) in addition to TB, offering potential synergies in the research and implementation of new, more efficient regimens in co-endemic areas.

We successfully employed a mouse model of *M. ulcerans* disease and achieved comparable clinical and microbiological results in the control groups as in previous experiments with the MU1059AL strain (24, 25). The decline in swelling grade and CFU in untreated mice can be attributed to a transition into a stationary phase about 7-9 weeks post-infection. Although the autoluminescent strain used in this experiment was previously reported to be fully virulent, we cannot exclude the possibility that it is slightly less virulent than its wild-type parent rendering it more prone to spontaneous clearance from the footpads (25).

In summary, high-dose RIF and RPT in combination with CLR performed well in a mouse model of *M. ulcerans* disease and warrant further investigation. These regimens can be administered orally; the elevated doses appear safe in humans when administered for 2 months or less (12, 13, 19). As BU has already been treated with rifampin and clarithromycin for several years, logistics and handling knowledge is already in place with health-care providers and a dose-adjustment could be easily implemented if shown effective in humans. One potential issue is that higher doses of rifamycins are likely to have a greater inductive effect on CLR metabolism in patients and could lead to subtherapeutic CLR exposures. We are currently evaluating other potential companion drugs for high-dose rifamycin regimens and will proceed to test whether they can cure *M. ulcerans* as short-course regimens in mice and ultimately, human patients.

## Acknowledgements

This study was supported by the National Institutes of Health (R01-AI113266). TO was supported by a personal grant from the Junior Scientific Masterclass at the University of Groningen, the Netherlands.

